# Random-effects meta-analysis of effect sizes as a unified framework for gene set analysis

**DOI:** 10.1101/2022.06.06.494956

**Authors:** Mohammad A. Makrooni, Dónal O’Shea, Paul Geeleher, Cathal Seoighe

## Abstract

Gene set analysis (GSA) remains a common step in genome-scale studies because it can reveal insights that are not apparent from results obtained for individual genes. Many different computational tools are applied for GSA, which may be sensitive to different types of signals; however, most methods test whether there are differences in the distribution of the effect of some experimental condition between genes in gene sets of interest. We have developed a unifying framework for GSA that first fits effect size distributions, and then tests for differences in these distributions between gene sets. These differences can be in the proportions of genes that are perturbed or in the sign or size of the effects. Inspired by statistical meta-analysis, we take into account the uncertainty in effect size estimates to reduce the influence of genes with greater uncertainty in effect size estimate on distribution parameters. We demonstrate, using simulation and by application to real data, that this approach provides significant gains in performance over existing methods. Furthermore, the statistical tests carried out are defined in terms of effect sizes, rather than the results of prior statistical tests measuring these changes, which leads to improved interpretability and greater robustness to variation in sample sizes. We also show that the approach naturally suggests alternative test types that are not usually considered in GSA; it can, for example, be applied to identify differences in effect size distributions between sample subgroups in a gene set of interest. Applying this approach to an analysis of gene expression changes between matched colon tumour and normal samples, we found several gene sets that showed distinct behaviour in patient subgroups with different prognoses. These may help to explain the clinical differences that have been reported between these patient groups.

**Author summary:** The role of gene set analysis is to identify groups of genes that are perturbed in a genomics experiment. There are many tools available for this task and they do not all test for the same types of changes. Here we propose a new way to carry out gene set analysis that involves first working out the distribution of the group effect in the gene set and then comparing this distribution to the equivalent distribution in other genes. Tests performed by existing tools for gene set analysis can be related to different comparisons in these distributions of group effects. A unified framework for gene set analysis provides for more explicit null hypotheses against which to test sets of genes for different types of responses to the experimental conditions. These results are more interpretable, because the group effect distributions can be compared visually, providing an indication of how the experimental effect differs between the gene sets. We can also apply this method to identify sets of genes that behave atypically in subgroups of samples. This enabled us to identify differences in the expression of several gene sets in colon cancer samples between individuals with reduced mortality and those without this benefit.

## Introduction

Gene set analysis (GSA) methods are used to provide insight into gene expression (or other genomics data types) by testing hypotheses on pre-defined sets of genes. This serves to leverage prior biological knowledge, reduce the number of hypotheses tested and improve the interpretability of the results [1]. Many different GSA methods exist [1], and they can be classified as either competitive or self-contained, based on the null hypothesis being tested [2]. When applied to gene expression data, for example, competitive methods test for enrichment of differentially expressed (DE) genes in gene sets relative to the background. Self-contained methods, on the other hand, assess whether gene sets contains DE genes, without comparing the extent of differential expression to background genes. Methods can be further categorized based on the direction of the expression changes that are the basis of the test. A directional hypothesis involves testing for either up-regulation or down-regulation of genes in a set, while a mixed hypothesis tests for differential expression regardless of direction [3].

The nature of the hypothesized difference between the inset and outset provides further method distinctions. For example, in over-representation analysis (ORA) [4] the null hypothesis is that the *proportion* of DE genes in the inset is not greater than the proportion in the outset. Despite being conceptually simple, ORA has been found to be the top performing commonly-used method in a review of GSA benchmarking and simulation studies [5]. However, by making a binary classification of genes as either DE or non-DE, based on an arbitrary p-value threshold, ORA discards information about the extent of the difference in expression and is tied to the power of the specific experiment performed. Instead of making a binary classification, GSEA [6] ranks genes by a ranking metric and then tests against a null hypothesis under which the rank distribution of a gene set is associated with group membership. Various ranking metrics have been used in many similar methods [1]. These methods have the advantage over ORA of taking the full gene rank into account, which reflects the size of the DE effect. Although metrics can be used that are influenced by both the effect size estimate and its uncertainty (such as the signal-to-noise ratio option in GSEA [7]), these must be collapsed into a single value for ranking purposes, resulting in a loss of information. The use of the signal-to-noise ratio in GSEA can also create difficulties for the inclusion of covariates and interaction terms in the experimental design [8]. Failure to account for these covariates could lead to spurious results in some cases and reduced ability to detect true enrichment in others. Ranking-based methods typically evaluate the significance of the differences between gene sets through sample permutation. This has the benefit of making the methods robust to failures in the assumptions (such as gene-gene independence) that are made by other methods, but at the expense of being computationally expensive and not suited to experiments with low sample numbers [9].

Here, we propose a novel approach to GSA that both provides a unifying framework for the different approaches outlined above and also takes into account the uncertainty in the estimate of the effect size from the first stage of the analysis. In our approach, the log fold change (LFC) for genes in a given set is modeled as a mixture of Gaussian distributions, with distinct components corresponding to up-regulated, down-regulated and non-DE genes. We use the Expectation Maximization (EM) algorithm to estimate the parameters of this mixture distribution. Using a methodology inspired by statistical meta-analysis [10], the standard error of the DE effect size estimate is incorporated into the estimation procedure, with genes with large standard errors having less influence on the parameter estimates than genes for which the DE effect is estimated with greater precision. A wide range of tests that are relevant for gene set analysis can be performed by applying model comparison techniques to estimated effect size distributions in different gene sets. We evaluated the performance of our approach relative to existing GSA implementations using both simulated data as well as real data derived from the The Cancer Genome Atlas (TCGA). Our method showed substantially increased power compared to existing methods in the simulations. When applied to real cancer data, our method recovered gene sets previously found to be significantly enriched for DE genes in different cancer types as well as highlighting differences in the perturbation of specific gene sets between sample subgroups that may help to explain clinical differences between these subgroups of cancer patients.

## Results

### Model and implementation

We model the LFC of genes as a mixture of three Gaussian components, corresponding to non-DE, up-regulated and down-regulated genes and make use of random effects meta-analysis to incorporate the variance in the gene-level LFC estimates in parameter estimation (see Methods for details). The statistical tests at the gene set level consist of comparison of the fit of models in which the LFC distribution is the same (null hypothesis) or different (alternative hypothesis) for genes in the inset (the gene set of interest) and outset (the remaining genes in the background set). We evaluated two types of model comparison that correspond to different kinds of GSA. The first consists of comparing the fits of a model in which all parameters of the LFC mixture distribution are shared between genes in the inset and outset to a model that allows the proportion of DE genes to differ between the two sets. We refer to this as the one degree of freedom (1DF) test. It evaluates whether the proportion of DE genes differs between inset and outset and can be modified to test, specifically, whether the proportion of DE genes is higher in the inset than outset. Using confidence intervals on the estimated proportion of DE genes we can obtain a lower bound on the difference in the proportion of DE genes in the inset and outset. This provides a useful means of ranking gene sets that takes account of the size of the difference in DE gene proportions between the gene sets as well as the uncertainty in this difference. In the second kind of test evaluated, we compared the fit of a model in which all parameters of the mixture distribution are allowed to differ between genes in the inset and outset to a model in which all parameters are shared. This test (referred to as the six degree of freedom, or 6DF test) is sensitive to any differences in the mixture distribution between the inset and outset (e.g. a difference in the size of the DE effect even if the proportion of DE genes is the same). Details of all of the tests are provided in Methods.

### Performance on simulated data

We first used simulations to assess method performance relative to existing methods. In the simulations the group effect was assumed to be normally distributed in the inset and outset, but with a larger variance in the inset, resulting in a larger number of DE genes with LFC above the threshold shown (Fig 1A). This is deliberately far-removed from our model, which fits a mixture of three normal distributions. The simulations illustrate a key feature that distinguishes our approach to GSA from many existing methods. GSA methods designed to identify gene sets with a higher proportion of DE genes typically perform statistical tests on the proportions of genes falling below an arbitrary p-value threshold; however, this proportion will depend on the number of samples. Indeed, it is likely to be the case that all genes are DE to some extent between any two biologically distinct groups of samples, with this difference becoming statistically significant, given sufficient sample numbers. Instead, we define DE by setting a threshold on the absolute value of the LFC and our method compares proportions of genes for which the true (but unknown) absolute value of the LFC is above this threshold. Our method significantly outperformed ORA, GSEA, and SAFE [11] (Figs 1C and 1D) on the simulated data. GSEA had low power with low sample numbers, but as sample numbers increased the number of permutations possible with which to derive the p-value leads to an increase in power. The false positive rate for GSEA, SAFE and ORA all tended to increase with increasing sample numbers (S1 Fig).

**Fig 1.**
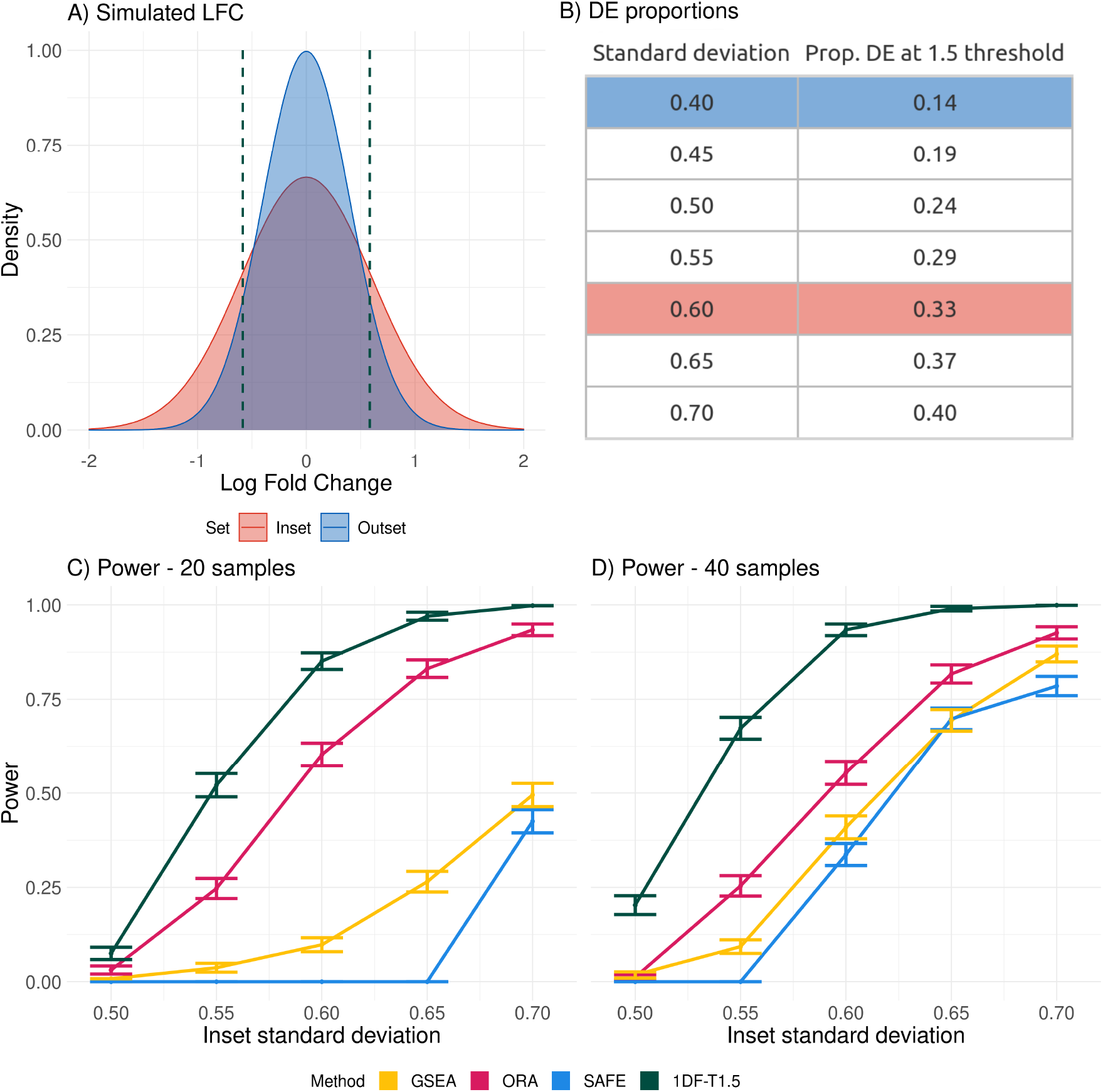
Power on simulated data. A) Simulated LFC distributions, with the 1.5 fold-change threshold shown as dashed lines. B) The proportion of DE genes in a gene set for each value of the standard deviation (SD) used. C) Comparison of the power of different methods applied to simulated data corresponding to 20 samples, as a function of different SD values for the LFC distribution in the inset (the SD in the outset remained fixed at 0.4). D) The power of the same methods when the number of samples was increased to 40.

Our method requires researchers to specify upfront the magnitude of the DE effects of interest. Unsurprisingly, the power of our method to detect the difference in the proportion of genes above the threshold in the simulated data depended on the relationship between the DE threshold and the LFC distributions in the inset and outset (Fig 2). The highest power is achieved when the threshold defines a region in the effect size distribution in which there is a substantial difference between the inset and outset. No such fold-change threshold is specified for existing enrichment-based methods (Figs 1C and 1D) and, instead, such methods often rely on p-value thresholds from statistical tests at the individual gene level. The lack of an explicit definition of differential expression in terms of the actual expression values in sample groups can lead to statistical inconsistencies. For example, the power of ORA can decrease with increasing sample numbers (Fig 1D). In figure 1D, this resulted from an increase in the power to detect the smaller expression changes in the outset as the sample number increases, causing a reduction in the difference in the proportion of genes detected as statistically significant. A threshold can also be applied in the original gene expression analysis when using ORA, with the test for enrichment applied to the proportion of genes that are statistically significant and have absolute LFC above the threshold. This resulted in an increase in power in ORA with increasing sample numbers (S4 Fig), highlighting the importance of explicitly defining the magnitude of the effects of interest.

**Fig 2.**
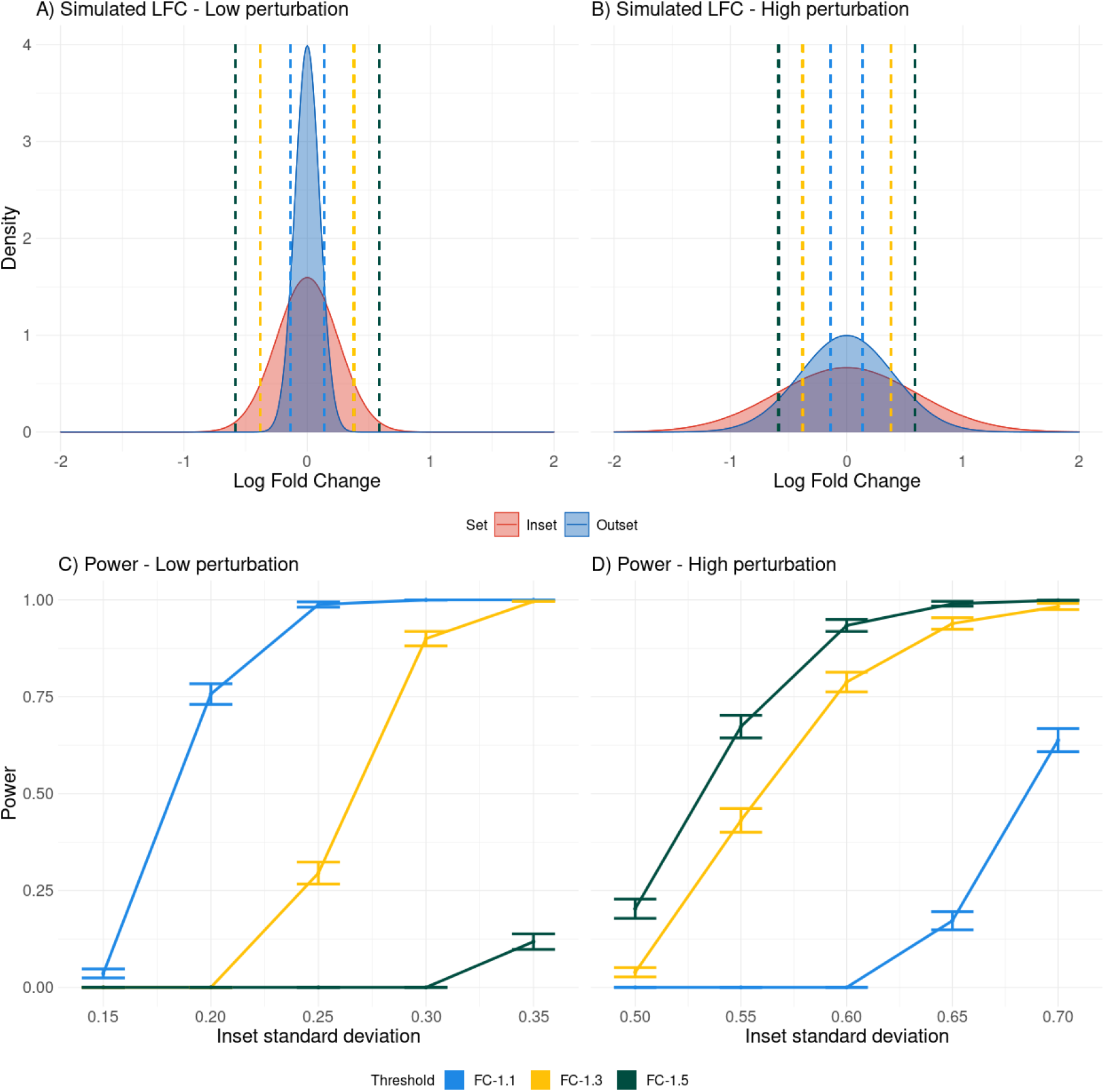
Different fold-change thresholds. A) Simulated LFC distributions with low LFC values. The thresholds used with our 1DF approach are shown as dashed lines. B) Simulated LFC distributions with high LFC values. C) The power of the 1DF test at different thresholds for 40 samples simulated with low LFC values. D) The power of the 1DF test at different thresholds for 40 samples with high LFC values.

### TCGA Data

We tested for enrichment of DE genes between tumour and matched normal samples from the TCGA database in 347 KEGG gene sets. The 1DF test identified 71, 51, and 51 gene sets as significantly enriched in TCGA-PRAD (prostate), TCGA-BLCA (bladder) and TCGA-COAD (colon), respectively, after correcting for multiple testing (S1 Table, S2 Table and S2 Table), and with a threshold set at a fold change of 1.5. This corresponds to approximately 20% of gene sets being called as significantly enriched in PRAD, and approximately 15% in both BLCA and COAD. Twenty-one gene sets were identified as significant in all three cancers when compared to matched normal samples (S4 Table). When ORA, GSEA, and SAFE [11], were applied on these datasets (with correction for multiple testing) several gene sets were identified as enriched in each cancer type (S5 Table, S6 Table and S7 Table), by ORA and the un-directional GSEA. SAFE failed to identify any significant gene sets supporting previous results which found SAFE to have low power [12]. To investigate the relationship between power and sample size for different GSA methods on real data, where the ground truth is unknown, we re-evaluated a gene set for which the evidence of enrichment was clear, as a function of decreasing sample numbers (Fig 3). The Calcium signaling pathway was identified as enriched by our method as well as by ORA, and GSEA; however, our method showed greater robustness to decreasing sample numbers. Consistent with the simulations, the behaviour of ORA as a function of the number of samples was erratic, although this may, in part, be explained by the fact that the gene set was only weakly significant with ORA in the full dataset. GSEA showed a consistent increase in power with increasing sample numbers although at low sample numbers the number of possible permutations is limited leading low power.

**Fig 3.**
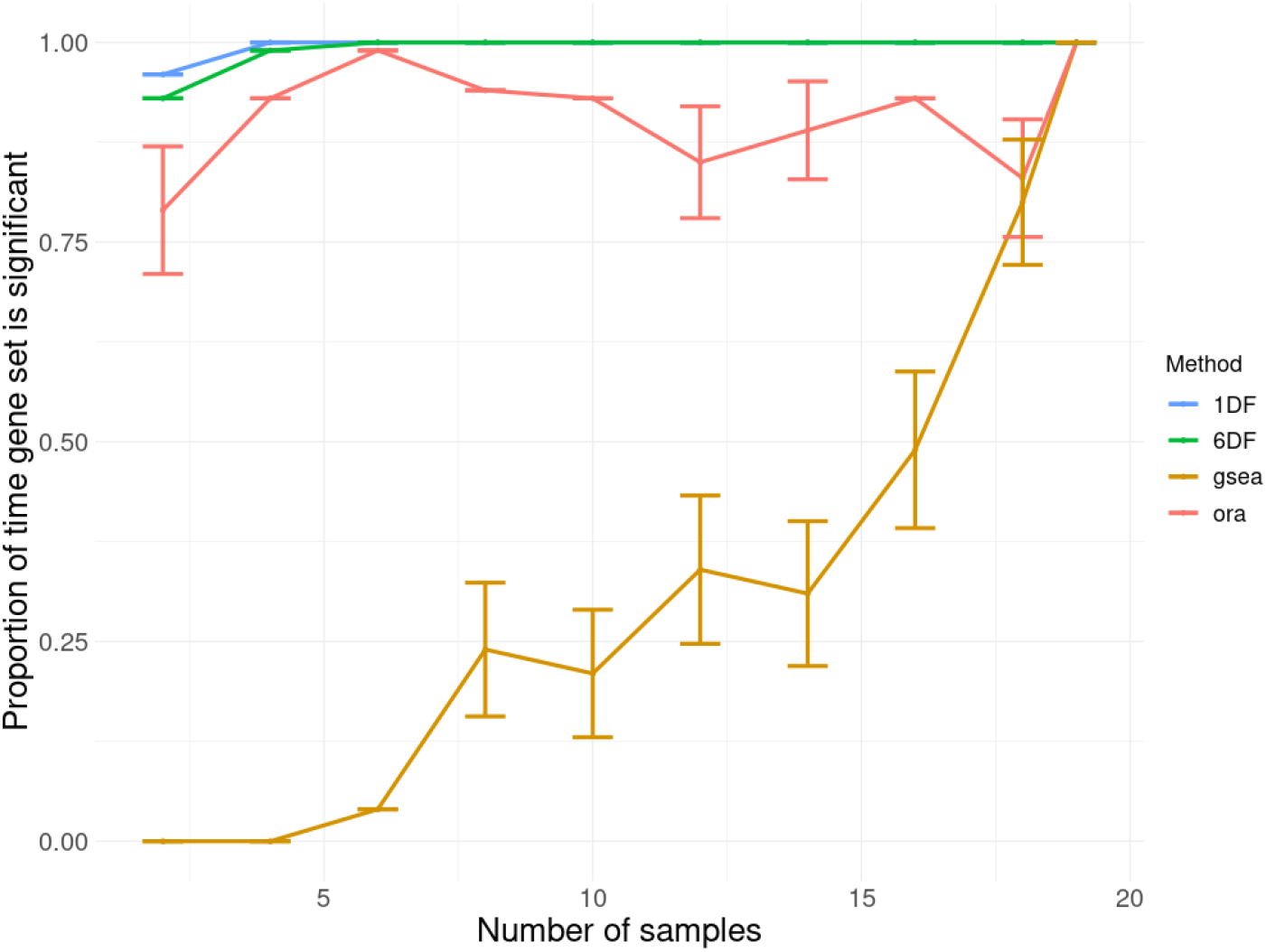
Power on down-sampled data. The proportion of times that the Calcium signalling pathway was identified as significant by each tool in TCGA-BLCA. For each sample there is matched-normal tissue and tumour tissue. For each sample number a random selection of samples and analysis was run 100 times.

Young females with colon cancer have reduced mortality compared to other colon cancer patients [13]. We applied our method to compare the LFC distribution associated with the group effect (cancer versus normal) between these and the remaining patients in order to investigate the distinctive features of colon tumours in young female patients. Considering all genes, the LFC distribution was significantly different in young females compared to the remaining samples (P = 9.2 × 10^-60^; 1DF test), with a substantially lower proportion of DE genes in the former. Fifty-nine individual gene sets had significant differences in the LFC between the patient groups, after correcting for multiple testing (S8 Table), fifty-five of which showed lower levels of DE in the young female subgroup. The fact that our statistical tests are based on comparisons of effect size distributions between sets of genes or sample subgroups has the advantage that the basis for the statistically significant differences can be visualized (Fig 4). We refer to the visualization of estimated effect size distributions in the inset and outset as post gene set analysis and our method may be helpful for this purpose even when the enriched gene sets are identified using another method.

**Fig 4.**
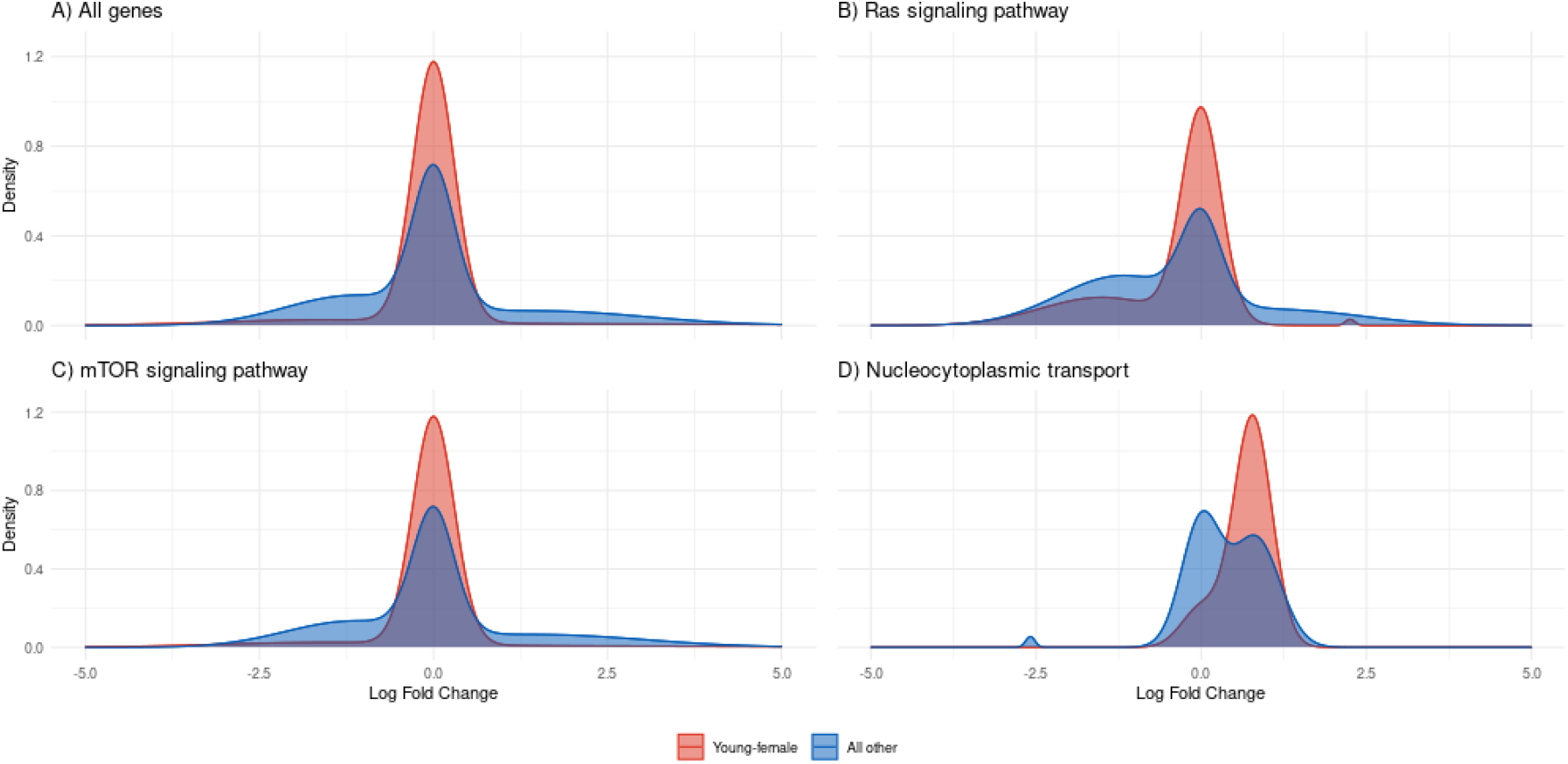
Estimated LFC distributions in COAD-TCGA. The LFC distribution, shown in red for the young female samples and in blue for all other samples, for all genes and for three gene sets of interest (see Discussion) that showed significant differences between the two groups of patients.

## Discussion

Many existing GSA methods test hypotheses that are defined in terms of the results of previous statistical tests, carried out on individual genes, rather than on the actual effect of interest. For example, the null hypothesis of enrichment-based methods (such as ORA) is that the proportion of genes passing the p-value significance threshold is the same in the inset and outset. This proportion depends on the power of the experiment and is liable to change in both sets if the experiment is repeated with a different number of samples, sometimes in unpredictable ways (Figs 1D and 3). Fundamentally, therefore, existing GSA methods provide results that relate to the specific experiment performed, rather than to the biological effects of interest. We propose an alternative approach to GSA in which the null hypotheses evaluated are expressed in terms of the underlying biology. Instead of asking, for example, whether the proportion of statistically significant genes differs between the inset and outset we assess whether the proportion of genes for which the true (but unknown) DE effect exceeds some specified threshold differs between the gene sets. This enables the researcher to consider the minimal effect size that may be of biological significance, rather than focusing on whether individual genes are statistically significant in a specific experiment.

The two tests we evaluated here consider distinct null hypotheses. The 1DF test evaluates whether the proportion of DE genes differs between the inset and outset, while the 6DF test is sensitive to differences in both the proportion of DE genes as well as the DE effect size. While the goal of the 1DF test is similar to that of existing enrichment-based GSA methods, the fold-change value at which one would consider a gene to be DE is not explicitly defined in ORA or in many other established GSA methods (although tools such as DESeq2 allow for LFC thresholds to be set when calling a gene as DE [14]). This is an important parameter that should be considered in the context of established methods as well, in order to reduce the influence of DE effects which, though statistically significant, are too small to be biologically meaningful. For example, in the extreme case of an infinite number of samples, there will be no error in the LFC estimates and all genes with non-zero fold-change will be statistically significant. Our approach involves setting a DE threshold in advance, enabling the researcher to define the effect sizes of interest and leading to more interpretable results. The null hypothesis for the 6DF test is that the LFC distributions for genes in the inset and outset are the same. This is a more general test that is sensitive to differences in both the proportion of affected genes and the magnitude of the effect. The results can be made more interpretable by visualizing the inferred effect size distribution (e.g. Figs 4, S2 Fig and S3 Fig), which often suggests why the null hypothesis is rejected for a specific gene set. Given GSEA makes full use of the rank distribution of genes, rejection of its null hypothesis can result from differences in proportions of DE genes or differences in the effect sizes of the DE genes or both. This is comparable to the 6DF test, though the reason for the rejection of the null hypothesis may be much less clear.

A further distinction between our approach to GSA and existing methods is that we take account of the uncertainty in the estimated group effect for each gene. Existing methods, by contrast, typically obtain a p-value or test statistic from an initial gene expression analysis and either count the number of genes with p-values below some threshold or or combine the uncertainty with the effect size in a ranking metric. By considering both the estimated effect and its standard error our method retains information about the size of the effect and the precision with which it has been estimated. This information is then used to estimate the parameters of a distribution describing the effect size in a set of genes, with the standard error of the estimate being used to reduce the influence on this distribution of effects that are poorly estimated. Taking account of this uncertainty resulted in an increase in power (S5 Fig). As expected, this improvement is greater for lower sample numbers where there is greater uncertainty in the effect size estimates. In this study we used a Gaussian mixture model (GMM) for the distribution of the logarithm of gene expression fold-change (LFC) between groups, because this allows the uncertainty in the effect size estimate to be incorporated easily into the estimation of the component variances within the M step of the EM algorithm. In principal, the GMM could be replaced with a more general model for the distribution of effects across genes, but this would require the development of efficient methods to incorporate the standard error of the effect sizes into the parameter estimation.

Our method can have a significant advantage over methods that are based on detecting differences in proportions of genes passing a p-value threshold using the hypergeometric distribution, particularly in the presence of relatively modest sample sizes or modest group effects that are shared across many genes in a gene set. This is because our method can accumulate evidence of a difference in the group effect distribution between gene sets, even when there is large uncertainty in the group effect for individual genes. In such a case, the proportion of DE genes passing the p-value threshold may be small, resulting in a low ability to detect enrichment of moderately DE genes. By contrast, where the DE status of a gene is clear, the advantage to avoiding hard assignment of genes as either DE or not DE is smaller. Consistent with this, our method retained power well with decreasing sample numbers (Fig. 3).

Methods, such as GSEA, that are based on sample permutation have the advantage of robustness, particularly to independence assumptions [7]. For example, ORA treats genes as independent observations, even though genes can share regulatory mechanisms, resulting in correlated expression across samples. This robustness comes at a cost, however, as sample permutation methods are computationally intensive and not useful for small sample numbers. The interpretation of results can also be challenging. The null hypothesis for GSEA is that the observed enrichment score for a gene set is not a result of differences between cases and controls. Rejection of this null hypothesis does not necessarily imply that genes within the gene set are affected disproportionately by the sample group, as the group effect is absent in the permuted samples. Furthermore the test statistic in GSEA is affected by both the proportion of DE genes and the magnitude of the effect and it is not obvious for significant gene sets whether a greater proportion of genes are perturbed in the gene set or whether the size of the perturbation is greater. Unlike ORA, which tests for a difference in the proportion of DE genes and the methods proposed here, which can test for a difference in proportions or for a difference in the effect-size distribution, GSEA does not distinguish the nature of the difference between the gene set and the background set. Our approach can also be adapted to use sample permutation, leading to greater robustness, but maintaining an advantage in terms of interpretability. For example, a probabilistic estimate of the proportion of genes in a gene set with group effect above a specified threshold can be obtained by summing over the gene-specific effect-size estimates (treating each effect size estimate as a random variable). A permutation-based test for a difference in this proportion between genes in the inset and outset can then be obtained by shuffling the sample labels. However, rerunning the full DE analysis for each permutation, is very computationally expensive using existing DE analysis tools.

Our 1DF method found many more significantly enriched gene sets than established methods between cancer and normal samples (S5 Table, S6 Table and S7 Table). Twenty-one gene sets were found to be significantly enriched in all three cancers (S4 Table), including several signaling pathways and several secretion pathways. Several cancer related gene sets were found to be perturbed in each cancer type but no one set was found to be common to all three cancers. Ranking gene sets by the lower bound on the difference in the proportion of DE genes in the inset and outset avoids a bias towards large gene sets that occurs when ranking by p-value. If ranking was done by the weight of the non-DE component alone this would bias the results towards smaller gene sets. In bladder cancer, ranking by the lower bound of the difference in weight leads to the gene set associated with malaria, one of the smaller gene sets, being ranked in sixth place, whereas when ranked by p-value it is ranked fifteenth. Given the potential use of anti-malarial drugs to treat different types of cancer [15] and bladder cancer more specifically [16] this may be of interest. The increased importance placed on the smaller vitamin digestion and absorption gene set in colon cancer is consistent with previous results that found perturbation of gene expression in this set to be closely linked to colon cancer progression [17]. In prostate cancer the nicotine addiction gene set is ranked as the most significantly enriched gene set despite its small size, supporting previous research indicating that this set is strongly perturbed in prostate cancer [18].

Colon cancer shows sexual dimorphism, with females under the age of 44 having a significant survival advantage compared to males of similar age and to older males or females [13]. Among the colon cancer samples we analyzed, the young-female patient group showed significantly less change in gene expression between the tumour and normal samples, compared to other patients (Fig 4A). There were fifty-nine individual gene sets with a significant difference in DE effect between these patient groups (S8 Table), with fifty-five sets showing less DE in the young-female samples. Because there were only two patients in the young-female group, this result should be considered as suggestive. Our method is sensitive to weak signals shared across multiple genes resulting in the capacity to retain power for small sample numbers (Fig 4); however, given the small sample number, we cannot rule out that the differences are caused by factors, other than membership of the young-female group, that are shared between these samples. The RAS signaling pathway, which is a mediator of several characteristics of cancer [19], and the Signaling pathways regulating pluripotency of stem cells were significantly less dysregulated in the young-female patients (Fig 4B). The mTOR signalling pathway, which is associated with tumors and with regulating metabolic pathways [20], and the Metabolic pathway gene set itself, were also among those showing lower levels of perturbation in the young-female patients (Fig 4C). By contrast, the nucleocytoplasmic transport gene set was more up-regulated in the young-female patients (Fig 4D). This is interesting, given the link between nucleocytoplasmic transport and cancer progression and the evasion of apoptosis [21]; however, a recent pan-cancer analysis reported no correlation between the expression of the nuclear transport factor 2 gene and overall survival or disease free survival in colon cancer [22], suggesting that its effect may depend on the cancer type.

## Conclusion

We describe a unified framework for GSA that supports the formulation and testing of different types of hypotheses relating to how genes with a shared annotation respond to an experimental condition of interest. Our approach can also be used to carry out a post-hoc analysis of enriched gene sets, providing information on *how* the gene set compares to other genes and not just on *whether* the gene set is enriched for genes that are perturbed under the experimental condition. We propose an approach to GSA based on this unified framework that can evaluate the range of hypothesis tests implicit in established GSA methods. This approach can provide increased power relative to established methods. When applied to real data our approach highlighted several gene sets relating to molecular pathways known to be involved in multiple cancer types, as well as highlighting gene sets and molecular pathways that show different effects in different sample groups.

## Materials and methods

### Gaussian Mixture Model (GMM)

We model the LFC using a mixture of Gaussian distributions, one with a mean of zero, for non-DE genes, and two with positive and negative means to capture positive and negative DE genes respectively. The LFC distribution is then given by:

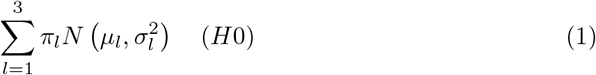

where *μ_l_*, 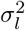 and *π_l_* are the mean, variance and weight, respectively, of component *l.* As a null hypothesis, we use a shared mixture distribution for all genes, regardless of gene set membership. In the alternative models gene set membership is included in the LFC distribution as follows:

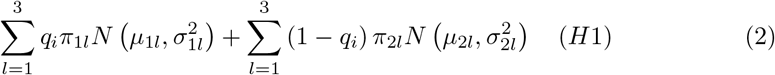

where *q_i_* is a binary value representing set membership. This allows the LFC distribution to differ between the inset and the outset. In the alternative model for the 1DF test, all genes contribute to the estimate of the means, variance and proportion of DE genes that are upregulated in the inset and the outset. The weight assigned to the non-DE genes in the inset is estimated using only genes in the set and the weight assigned to the non-DE genes in the outset is estimated using only genes outside the set. In the alternative model for the 6DF approach all parameters for the inset distribution are estimated using the genes in the set while all parameters in the outset distribution are estimated using genes outside the set.

Our method uses a soft threshold, *τ*, to define biologically relevant DE. We place the constraint on the parameters of the mixture components corresponding to upregulated and downregulated genes such that at most 0.25 of the area of either component falls in the interval [–*τ*, +*τ*]. The variance of the non-DE component is fixed such that 95% of the area of the component lies within this threshold region.

The 95% confidence interval for the non-DE component weight was used to rank significant gene sets, with genes ranked highest having the largest difference between the lower bound of this weight in the inset and its upper bound in the outset. The standard error for the estimate of the weight of the non-DE component (used in the calculation of the 95% confidence interval) was derived from the square root of the inverse of the hessian matrix of the negative log-likelihood function. This was calculated using the numDeriv package in R.

### Mixture Random-Effects Meta-analysis

The GMM described above does not consider the standard error associated with the LFC estimate for each gene. Each estimated LFC comes with uncertainty, in other words each estimated LFC is the sum of true LFC and an error term. We fit a GMM to the true LFC’s and we also assume that the error term is normally distributed with mean zero and standard deviation equal to the standard error of the estimated LFC (*σ_i_*, corresponding to each gene). Compounding these distributions we obtain the following LFC distribution:

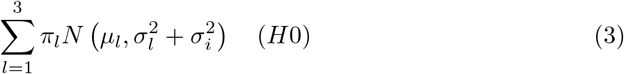

Similar to the previous section, we can build the alternative model by adding the binary indicator for the set membership:

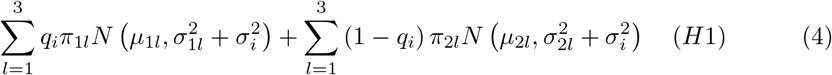

Considering each gene as a single study and interpreting 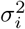 as the within study variance and 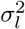 as between study variance, we see that our model here is similar to the standard random effect meta analysis except that the underlying distribution of LFC is a GMM instead of a single distribution. To estimate the parameter of the model we use Expectation Maximization algorithm. In the M-step, we use an iterative process (between the mean and variance of each component in the mixture of Gaussian) as unlike the GMM, the contribution of each gene to the parameters is weighted not only by the posterior probability (calculated in the E-step) but also by a weight whose formula is given by (following [10]):

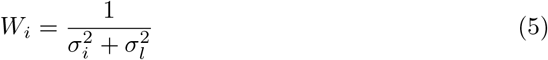

### Simulations

We simulated RNA-Seq gene expression data for *N* individuals in two equally sized groups. The genes were divided into S non-overlapping gene sets of equal size. We sampled gene expression counts for each gene, *i,* from the negative binomial distribution *NB*(*μ_i_, φ_i_*). In order to choose realistic parameters for this distribution, we used a real data set. The mean and the dispersion parameters for all genes in this data set were estimated using DESeq2 and the BRCA-TCGA data. For each gene in the count data matrix, we randomly chose a mean and a dispersion parameter from these values. The expression values for non-DE genes were again sampled from *NB*(*μ_i_, φ_i_*) and expression values for DE genes were sampled from *NB*(*μ_i_, φ_i_*) for one sample group and from *NB*(*FCμ_i_, φ_i_*) for the other, with *FC* the fold change value. Equal numbers of DE genes were up- and down-regulated.

LFC values for the inset and the outset were drawn from a normal distribution with a mean of zero and the standard deviation differing between the inset and the outset. In each power simulation the enriched inset of 100 genes was compared to an outset of 9,900 genes and repeated 1,000 times. In order to correct for multiple hypothesis testing p-values were obtained for 9,000 non-enriched sets and combined with the p-values from the power simulations and all p-values were corrected using the Benjamini-Hochberg procedure. This mirrors the approach used by Ma *et al.* (2020) [23]. Using DESeq2 [14] we obtained an estimate of the LFC and the standard error of this estimate. Established methods where implement using the *EnrichmentBrowser* package in R [24], with the exception of GSEA which was run using the absolute signal to noise ratio as the ranking metric [25].

### Real Data

Gene counts and clinical data for the TCGA datasets were retrieved through the GDC portal. The KEGG gene sets were retrieved using the GSEABenchmarker package [3]. For TCGA-BLCA there were 19 matched tumour and normal samples, for TCGA-PRAD, 51 matched tumour and normal samples and for TCGA-COAD, 41 matched tumour and normal samples. When comparing the menstruating and non-menstruating patients with colon cancer there were two menstruating patients and 36 non-menstruating patients each with a tumour and a normal sample. Lowly expressed genes were removed before differential gene expression analysis using edgeR with the default parameters [26]. DESeq2 was used again to obtain a LFC estimate and the standard error of this estimate, with the patient information included in the design formula. When comparing menstruating and non-menstruating patients, group membership was included as an interaction term.

## Supporting information

**S1 Fig Simulation FPR** False positive rate by sample number and by level of DE.

**S2 Fig Distribution estimate - Low variance** Estimates of the LFC distribution of the inset.

**S3 Fig Distribution estimate - High variance** Estimates of the LFC distribution of the inset.

**S4 Fig Simulation power - ORA** Power of ORA at different LFC thresholds.

**S5 Fig GMM with random effects vs GMM only** The improvement observed with the inclusion of the random-effects in our approach.

**S1 Table TCGA-PRAD significant sets** Gene sets identified by 1DF test with a threshold of 1.5.

**S2 Table TCGA-BLCA significant sets** Gene sets identified by 1DF test with a threshold of 1.5.

**S3 Table TCGA-COAD significant sets** Gene sets identified by 1DF test with a threshold of 1.5.

**S4 Table Significant sets across cancers.** Gene sets identified as enriched in three cancer types.

**S5 Table TCGA-PRAD significant sets** Gene sets identified by other tools.

**S6 Table TCGA-BLCA significant sets** Gene sets identified by other tools.

**S7 Table TCGA-COAD significant sets** Gene sets identified by other tools.

**S8 Table Direct comparison** – Significantly different KEGG gene sets between sample subsets in COAD.

There are no primary data in the paper; all materials are available at https://github.com/osedo/GSA-MREMA and we have archived our code on Zenodo ()

## Acknowledgments

This publication has emanated from research conducted with the financial support of Science Foundation Ireland under Grant number 16/IA/4612. DOS was funded through Science Foundation Ireland Grant Number 18/CRT/6214. Paul Geeleher is supported by the NIH, including a K99/R00 award from NHGRI (5R00HG009679-03) and an R35 award from NIGMS (1R35GM138293-01). The results published or shown here are in whole or part based upon data generated by the TCGA Research Network: https://www.cancer.gov/tcga.

